# Structural dynamics of E6AP E3 ligase HECT domain and involvement of flexible hinge loop in ubiquitin chain synthesis mechanism

**DOI:** 10.1101/2022.11.18.516873

**Authors:** Kazusa Takeda, Ikumi Muro, Fuminori Kobayashi, Holger Flechsig, Noriyuki Kodera, Toshio Ando, Hiroki Konno

**Author notes:** These authors contributed equally to this work (Kazusa Takeda, Ikumi Muro). Correspondence and request for materials should be addressed to HK.

## Abstract

Ubiquitin (Ub) ligases E3 are an important factor in selecting target proteins for ubiquitination and determining the type of polyubiquitin chains on the target proteins. In the HECT (homologous to E6AP C-terminus)-type E3 ligases, the HECT domain is composed of an N-lobe containing the E2-binding site and a C-lobe containing the catalytic Cys residue that forms a thioester bond with Ub. These two lobes are connected by a flexible hinge loop. The large conformational rearrangement of the HECT domain via the flexible hinge loop is essential for HECT-type E3-mediated Ub transfer from E2 to a target protein. However, detailed insights into the structural dynamics of this type of E3 ligases remain unclear. Here, we provide the first direct demonstration of structural dynamics of the E6AP HECT domain using high-speed atomic force microscopy. We also investigated structural dynamics of hinge loop flexibility restricted HECT domain, and we found that flexibility of the E6AP hinge loop has a great impact not only on its structural dynamics but also on the formation of free Ub chains mediated by E3-E3 interactions.

## Introduction

Ubiquitin (Ub) is a 76 amino acid protein highly conserved in eukaryotic organisms (1). Ub conjugation to target proteins (ubiquitination) is a post-translational modification that regulates numerous cellular processes (2). Ubiquitination of target proteins is accomplished by isopeptide bond formation between the carboxy group of the C-terminal glycine (Gly) residue of Ub and the ε-amino group of lysine (Lys) side chains on target proteins (3). The formation of an isopeptide bond between Ubs that gives rise to a poly-Ub chain on the target proteins. The types of poly-Ub chains depends on which of the seven Lys residues or the N-terminal methionine (Met) residue (M1, K6, K11, K27, K29, K33, K46, K63) on Ub is used for chain elongation (4). Ub conjugation and poly-Ub chain elongation on a target protein are accomplished by a series of enzymes, including a Ub-activating enzyme (E1), a Ub-conjugating enzyme (E2), and a Ub ligase (E3) (5). The K48 poly-Ub chain leads to proteolysis of the substrate (6), the K63 poly-Ub chain is involved in DNA repair and signal transduction (7, 8), and the linear M1 poly-Ub chain is involved in the nuclear factor-kappa B activity (9).

E3 is the most important factor in selecting a target substrate proteins and determining the type of Ub chain (10). Depending on the difference in the ubiquitination mechanism, E3 proteins can be largely classified into two: the homologous to E6AP C-terminus (HECT)-type and the really interesting new gene (RING)-type. RING-type E3 functions as a scaffold and adds Ub directly from E2 to a substrate protein (11). In contrast, HECT-type E3 forms a thioester bond between the carboxyl group of the C-terminal Gly of Ub and the conserved Cys residue on the HECT domain (see below) of the E3 ligases before transferring Ub to a target protein (12).

The diverse N-terminus regions of HECT-type E3 ligases mediate target protein recognition. Based on the N-terminal architecture, HECT-type E3 ligases have been classified into three groups: the NEDD4 family containing WW domain, the HERC (HECT and RCC1-like domain) family containing regulator of chromosome condensation 1 (RCC1)-like domains (RDLs), and others (12). The C-terminus of approximately 350 amino acids containing a catalytic Cys residue is defined as the HECT domain and is well conserved in all HECT-type E3 ligases. The HECT domain is composed of an N-lobe containing the E2-binding site and a C-lobe containing the conserved catalytic Cys residue (13). These lobes are connected by a flexible hinge loop. The introduction of a mutation that reduces the flexibility of the hinge loop results in the loss of ubiquitin transfer to target proteins (14). These observations led to the hypothesis that a large conformational rearrangement of the HECT domain via the flexible hinge loop is essential for Ub transfer by HECT-type E3 ligase. In addition, the HECT domain synthesizes the unanchored poly-Ub chains, and the Ub_2_ chain is the principal product (15–17). Although the physiological relevance of the Ub2 chain still remains unclear, understanding its formation mechanism is useful to gain an insight into the mechanism underlying Ub transfer from E2 to E3. Detailed information of the structural dynamics of HECT-type E3 ligases is also essential for a more comprehensive understanding of their functional mechanism.

To this end, here we used high-speed atomic force microscopy (HS-AFM) to investigate the structural dynamics of the HECT domain of E6AP E3 ligase. The orientation of the C-lobe relative to the N-lobe in the HECT domain was determined and compared to previous structural studies. We also determined the structural dynamics of a mutated HECT domain in which the flexibility of the hinge loop was restricted by the substitution of GSRN with PPPP. Biochemical analyses also determined effects of this mutation on ubiquitination and Ub2 production, and elucidated the underlying mechanisms. Based on these results, we propose structural changes occurring in E3 during Ub transfer.

## Results

### HS-AFM observations of the E6AP HECT domain

The crystal structure of a complex between the E6AP HECT domain and the UbcH7 E2 enzyme (PDB, 1C4Z) showed that the catalytic Cys of the E2 is separated by 4.1 nm from the catalytic Cys of the HECT domain (Fig. 1A). In this structure, the HECT domain is L-shaped as also seen in the crystal structure of the HECT domain alone (PDB, 15DF) (Fig. 1B, left). This distance is too far for Ub transfer. This distance is shortened to 1.6 nm, when the HECT domain is T-shaped as found in the crystal structure of the WWP1 HECT domain alone (Fig. 1B, middle) but the distance is still too far for Ub transfer (13, 14). The distance is significantly shortened to ~ 0.8 nm in the structure of the HECT domain of neuronal precursor cell-expressed developmentally downregulated 4 (NEDD4) (catalytic conformation state), allowing transfer of Ub from E2 to the C-lobe (5) (Fig. 1B, right). Therefore, a large conformational change of the C-lobe via the flexible hinge loop enables Ub transfer from E2 to HECT-type E3 (5, 13, 14). The HS-AFM observation of the wild-type HECT domain of E6AP (E6AP^HECT_WT^) exhibited two distinct particles: spherical and oval ones (Fig. 1C). As described above, previous studies of the crystal structures of the HECT domain without Ub have revealed three structural states: L-shape, reversed T-shape, and the catalytic conformation (Fig. 1B). To assess the structural state of each particles appeared in the HS-AFM images, we constructed simulated AFM images from the crystal structures of E6AP complexed with UbcH7 (PDB, 1D5F), WW domain containing E3 Ub protein ligase 1 (WWP1) (PDB, 1ND7), and NEDD4 (PDB, 3JVZ) (Fig. 1D, Figs. S1 and S2). By comparison between real (Fig. 1C) and simulated (Fig.1D) AFM images, the oval particles are considered to correspond to the L-shape, while the spherical particles correspond to the reversed T-shape or the catalytic conformation. Because the appearances of simulated AFM images of the reversed T-shape and the catalytic conformation were similar, distinguishing these conformations was difficult. The successive HS-AFM images captured at 6.7 frames per sec (fps) showed dynamic transitions between the oval and spherical appearances in the same single molecules (Fig. 1E). The transitions occurred at the timescale of ~170 ms (Fig. 1F). We further attempted to distinguish conformational states of individual imaged particles by their height. The height of spherical particles (4.1 nm on average) was only slightly higher than oval particles (3.8 nm on average), as shown in Fig. 1G. However, the simulated AFM images showed approximately the same height among the three conformational states (Fig. 1D), disabling state identification by height measurements. Nevertheless, the HS-AFM imaging of E6AP^HECT_WT^ could distinguish between L-shape and reversed T-shape/catalytic conformation and detect dynamic transitions between the two conformational groups. Next, we carried out HS-AFM imaging of E6AP^HECT_WT^ containing Ub (Fig. 2A). Unlike the structure of the HECT domain alone (without Ub), small particles appeared around the bulky HECT domain. These small particles are Ub. Even in the Ub-containing sample, the HECT domain appeared as interchangeable spherical and oval particles (Fig. 2B–D). This observation indicates that the HECT domain can adopt both structural states, irrespective of the absence and presence of Ub. We estimated the dwell time in each structural state in the presence of Ub. In contrast to the HECT domain without Ub (Fig. 1F), there was a difference in the dwell time between the two structural states; the dwell time of oval shape was longer (212 ms) than that of spherical shape (138 ms) (Fig. 2E). This indicates that HECT domain tends to take L-shape after receiving Ub. A previous biochemical study estimated that the entire Ub transfer,E1 → E2(UbcH7) → E3(E6AP) → target protein (p53), takes place approximately in 0.5 min (18). Therefore, the conformational transitions of the HECT domain between spherical and oval shapes dynamically occur many times during the entire Ub transfer process, and therefore, should not be rate-limiting events in the transfer process.

**Fig. 1.**
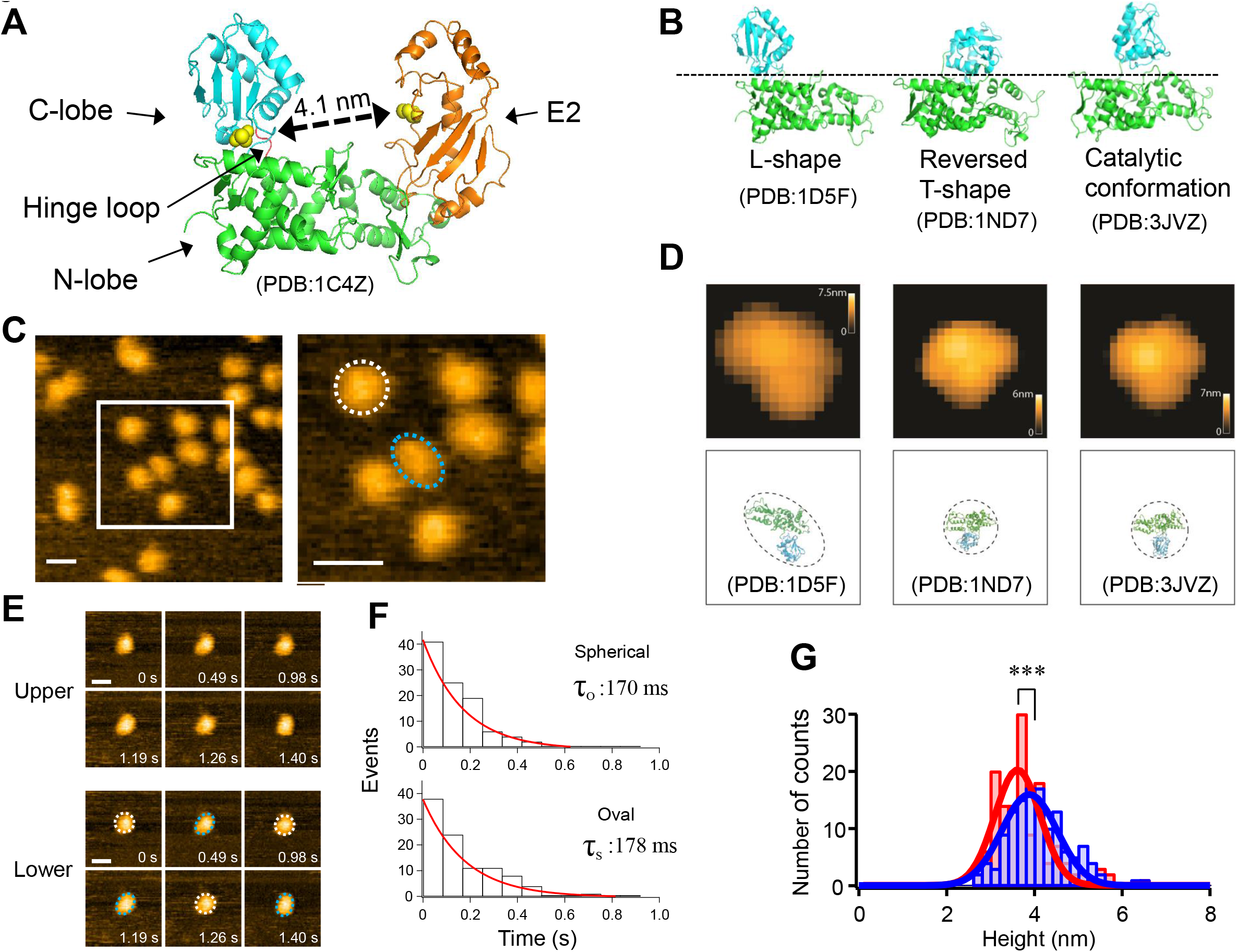
Three different structural states of HECT domain and HS-AFM imaging of E6AP^HECT_WT^. (A) Structure of an E6AP HECT domain-E2 complex. The HECT domain is composed of N-lobe (green) and C-lobe (cyan), which are connected by a hinge loop (red). The ubiquitin-conjugating enzyme (E2) is shown in orange. The Cys residue on the HECT domain and E2 that form thioester bonds with Ub are shown as yellow spheres. The distance of each Cys residue between the HECT domain and E2 is about 4.1 nm, and a large conformational change of the C-lobe is required to pass Ub. (B) Left, the structures of E6AP HECT domain showed that the catalytic Cys residue of E2 is separated from that of the E6AP HECT domain by 4.1 nm as described in (A) (L-shape), which is too far for Ub transfer to be possible. Middle, the Cys residue of E2 is located within 1.6 nm of that of WWP1 HECT domain (reversed T-shape) but the distance is still far for Ub transfer (13, 14). Right, the distance is significantly shortened (~ 0.8 nm) in the structure of NEDD4L HECT domain (catalytic conformation state) and it can be transfer Ub from E2 to C-lobe. (C) Left, AFM imaging of E6AP^HECT_WT^. The operational parameters were scan range, 100 × 100 nm (100 × 100 pixels); scan rate, 150 ms/frame; and z-scale, 4 nm, bar 15 nm. Right, magnified image of the region surrounded by the white line in the left panel. The typical two different shape of particles, spherical (dotted white line) and oval (dotted blue line), are indicated. (D) Automatized fitting of PDB structures into experimental AFM images (see Materials and Methods for details, and Figs. S1 and S2). The upper panel shows simulation AFM images of molecular orientations identified from fitting corresponding PDB structures displayed in the lower panel to HS-AFM images (molecular structures are displayed in scale). The image correlation coefficient of oval shape toward L-shape (PDB:1D5F) was 0.96. The image correlation coefficient of spherical shape toward reversed T-shape (PDB:1ND7) and catalytic conformation (PDB:3JVZ) were 0.95 and 0.94, respectively. The maximum heights were 5.4 nm for L-shape, 4.9 nm for reversed T-shape and 5.9 nm for catalytic conformation, respectively. (E) Upper, HS-AFM image sequence of E6AP^HECT_WT^. Lower, the same HS-AFM image sequence of E6AP^HECT_PPPP^ shown in the upper panel. The spherical and oval shapes of the HECT domain in each frame is denoted by the dotted white lines and the dotted blue lines, respectively. The inter conversion of the structure between spherical (dotted white lines) and oval particles (dotted blue lines) were observed. Scan range, 40 × 40 nm (60 × 60 pixels); scan rate, 70 ms/frame; z-scale, 4 nm, bar 10 nm. (F) Dwell time distributions in oval and spherical shape of E6AP^HECT_WT^. Distributions are fit to a single exponential decay to obtain the time constants (τ_O_ and τ_S_). (G) Height distributions of the oval (red) and the spherical (blue) particle. The corresponding solid lines are most probable Gaussian fitting obtained with 3.8 ± 0.7 nm (oval particle) and 4.1 ± 0.7 nm (spherical particle) for mean height ± S.D. The mean height in spherical particle was compared with the height in oval particle: ***p < 0.001; note that significant difference was observed between the oval particle and the spherical particle. The sample numbers are 139 in oval particle, 129 in spherical particle, respectively.

**Fig. 2.**
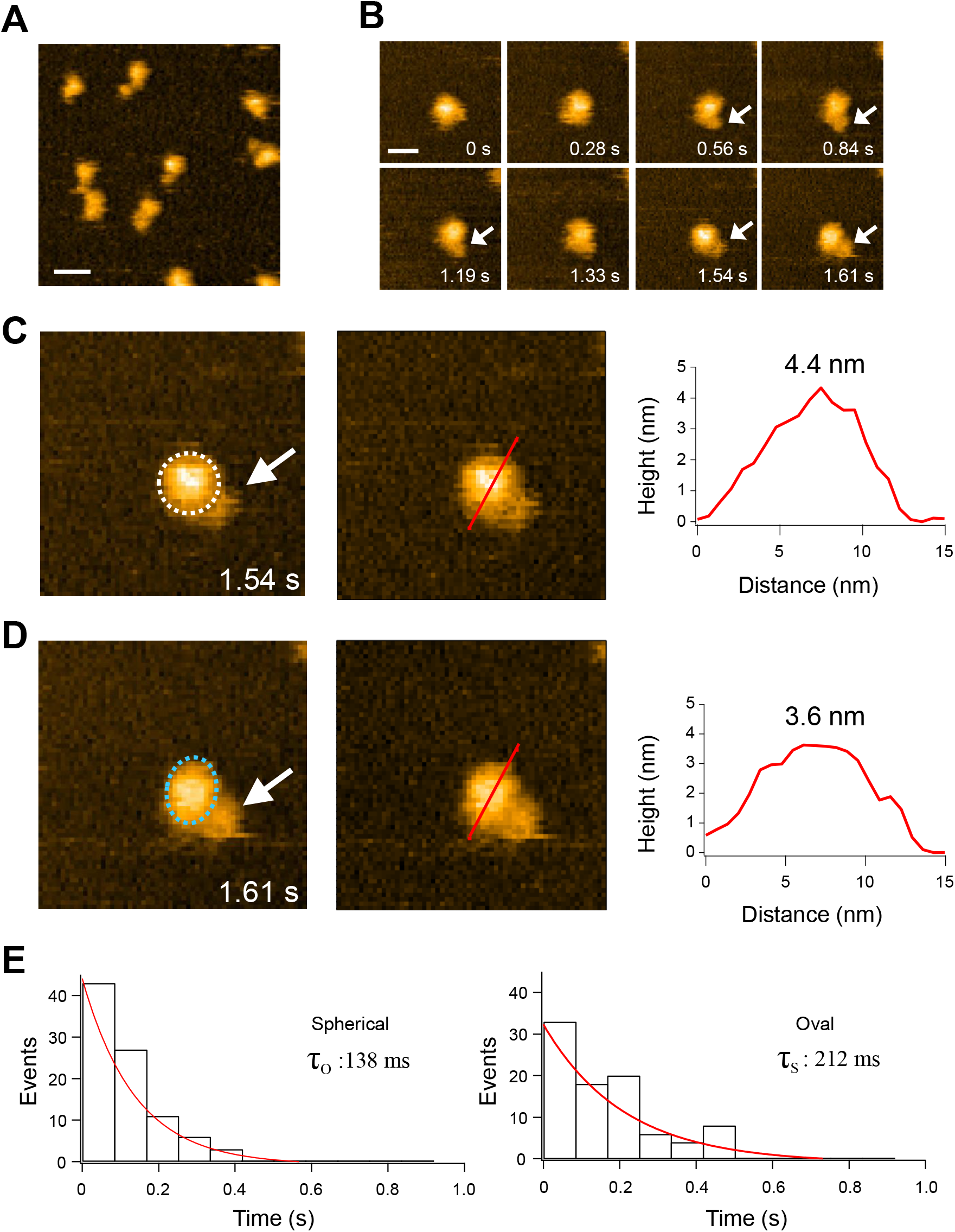
HS-AFM imaging of E6AP^HECT_WT^ containing Ub. (A) Representative HS-AFM image of E6AP^HECT_WT^ containing Ub. Scan range, 100 × 100 nm (100 × 100 pixels); scan rate, 150 ms/frame; z-scale, 4 nm, bar 15 nm. (B) HS-AFM image sequence of E6AP^HECT_WT^ containing Ub. Scan range, 40 × 40 nm (60 × 60 pixels); scan rate, 70 ms/frame; z-scale, 4 nm, bar 10 nm. The white arrows denote the small particles around the HECT domain, which are apparently Ub. (C) Left, the same images acquired at 1.54 s in (B). The shape of the HECT domain, spherical shape, at 1.54 s is indicated by the dotted white lines. The white arrows denote the small particles around the HECT domain, which are apparently Ub. Middle, height measurement of the spherical shape in the left panel. Right, line profile of the red line in the middle panel. The highest height in the line profile, 4.4 nm, is also shown. (D) Left, the same images acquired at 1.61 s in (B). The shape of the HECT domain, oval shape, at 1.61 s is indicated by the dotted white lines. The white arrows denote the small particles around the HECT domain, which are apparently Ub. Middle, height measurement of the oval shape in left panel. Right, line profile of the red line in middle panel. The highest height in the line profile, 3.6 nm, is also shown. (E) Dwell time distributions in oval and spherical shape of E6A^PHECT_WT^ containing Ub. Distributions are fit to a single exponential decay to obtain the time constants (τ_O_ and τ_S_).

### Structural state of mutant HECT domain (E6AP^HECT_PPPP^) lacking hinge loop flexibility

Next, to clarify the effect of the flexible hinge loop on the conformational transitions of the HECT domain, we carried out HS-AFM imaging of a mutated HECT domain with a much less flexible hinge. A previous study has already clarified that the flexibility in the hinge loop is lost by deletion of the hinge loop or substitution of the amino acids on the hinge loop to proline. (14). Therefore, we prepared a mutant HECT domain (E6AP^HECT_PPPP^) lacking hinge loop flexibility by replacing the amino acids of the hinge loop with proline (Fig. 3A and 3B). The HS-AFM images of E6AP^HECT_PPPP^ showed only spherical particles; oval particles were not observed and therefore no structural transitions occurred (Figs. 3C & 3E), indicating that the substitution of proline has indeed lost flexibility in the hinge loop. However, the mean height of the spherical particles was 3.7 nm, close to the height of oval particles appeared in the E6AP^HECT_WT^ sample (Fig. 3D). Since the crystal structure of E6AP^HECT_PPPP^ is not available, we obtained a predicted 3D structure of E6AP^HECT_PPPP^ by using AlphaFold2 (19) (Fig. 3F, right). To check the reliability of this method, we also obtained a predicted structure of E6AP^HECT_WT^ in the same way. The predicted structure of E6AP^HECT_WT^ (Fig. 3E, left) was very similar to the structure with L-shape previously revealed by X-ray crystallography (PDB, 1D5F) (14). The predicted mutant structure E6AP^HECT_PPPP^ was much different from the L-shape and reversed T-shape conformations (RMSD of 11.4Å and 11.9Å, respectively), and instead was similar to the catalytic conformation (RMSD of 7.6Å). AFM simulations of the AlphaFold2 E6AP^HECT_PPPP^ structure (Fig. 3G) not only clearly confirm the spherical shape topography, but also indicate a lower height magnitude (~4nm) as compared to simulation AFM images of wild-type shapes (Fig. 1D), which supports the analysis based on HS-AFM (Fig. 3C). The obtained results evidence that the mutant HECT domain lacking hinge loop flexibility preferentially adopts a spherical shape (reversed T-shape or catalytic conformation state).

**Fig. 3.**
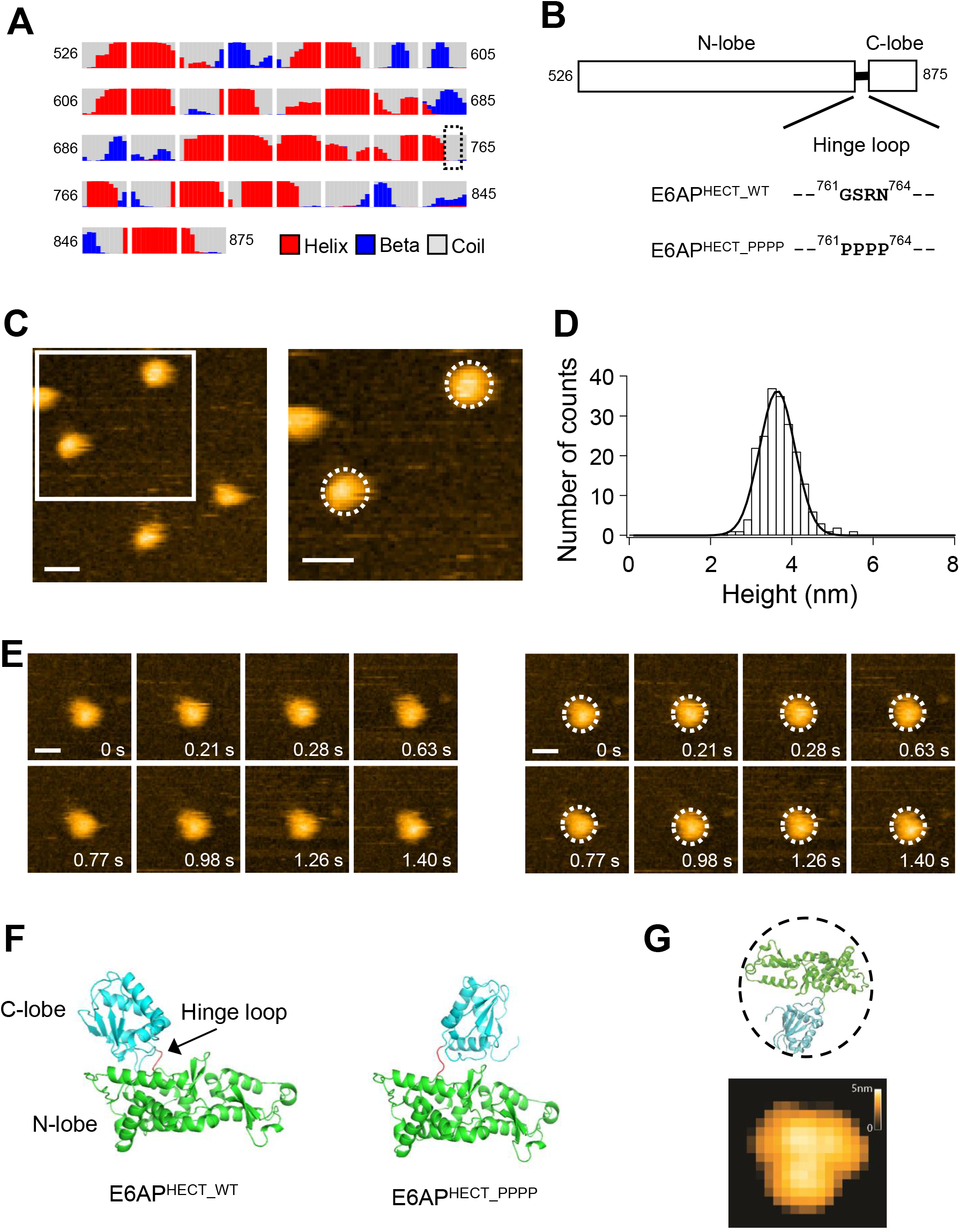
HS-AFM imaging and structure prediction of E6AP^HECT_PPPP^. (A) Secondary structure prediction of E6AP^HECT_WT^ by RaptorX property (http://raptorx.uchicago.edu/StructurePropertyPred/predict/). The percentages of predicted structural properties in each amino acid are shown in helix (red), *β*-strand (blue) and coil (gray). The amino acids tht constitutes hinge loop (GSRN) is boxed. (B) The amino acid sequence of the hinge loop flexibility restricted mutant HECT domain, E6AP^HECT_PPPP^. (C) Left, AFM image of E6AP^HECT_PPPP^, Scan range, 100 × 100 nm (100 × 100 pixels); scan rate, 150 ms/frame; z-scale, 5 nm, bar 15 nm. Right, magnified image of the region surrounded by the white line in the left panel. The typical spherical shapes of the particles are denoted by the dotted white line. (D) Height distributions of the E6AP^HECT_PPPP^. The corresponding solid lines are most probable Gaussian fitting obtained with 3.7 ± 0.5 nm for mean height ± S.D. The sample number is 200. (E) Left, HS-AFM image sequence of E6AP^HECT_PPPP^ from 0 s to 1.40 s. Scan range, 40 × 40 nm (60 × 60 pixels); scan rate, 70 ms/frame; z-scale, 4 nm, bar 10 nm. Right, the same HS-AFM image sequence of E6AP^HECT_PPPP^ shown in the left panel. The spherical shape of the HECT domain in each frame is denoted by the dotted white lines. (F) Structure prediction of E6AP^HECT_WT^ and E6AP^HECT_PPPP^ by AlphaFold2. Left, the result of structure prediction of E6AP^HECT_WT^. N-lobe, C-lobe and hinge loop of E6AP HECT domain are shown in green, cyan and red, respectively. Right, the result of structure prediction of E6AP^HECT_PPPP^. (G) AFM simulation of the E6AP^HECT_PPPP^ structure predicted by AlphaFold2. The upper panel shows the molecular structure of E6AP^HECT_PPPP^. The lower panel shows the corresponding simulation AFM image.

### Relationship between HECT domain hinge loop flexibility and Ub_2_ formation mechanism

While a E6AP mutant with an unflexible hinge loop does not deliver Ub to target proteins (14), it may be possible that E2 delivers Ub to the catalytic Cys residue on E6AP^HECT_PPPP^, although the hinge flexibility is lost. This is because HS-AFM observations revealed that the Cys residue on the C-lobe of E6AP^HECT_PPPP^ is located close to the N-lobe to which E2-Ub binds (Fig. 3). Since E6AP^HECT_WT^ is known to synthesize unanchored poly-Ub chains (mainly a Ub_2_ chain) (15–17), we performed biochemical experiments to examine whether E6AP^HECT_PPPP^ could also synthesize unanchored poly-Ub chains. This biochemical analysis possibly provides an insight into the mechanism of Ub transfer from E2 to E3. Remarkably, E6AP^HECT_PPPP^ yielded free Ub_2_ (neither bound to the E2 nor E6AP^HECT_PPPP^) with a significantly higher efficiency than E6AP^HECT_WT^ (Fig. 4A). It is reported that KIAA10-CD protein, a kind of HECT having a much higher Ub_2_ formation efficiency than E6AP, forms Ub_2_ by a different formation mechanism with E6AP (16). It is possible that the hinge loop of E6AP HECT may influence Ub_2_ formation mechanism. Therefore, we decided to investigate whether efficient Ub_2_ formation by E6AP^HECT_PPPP^ is due to a different Ub_2_ formation mechanism. So far, Wang et al. proposed three possible mechanisms for Ub_2_ chain synthesis using the HECT domain (Fig. 4B) (16). In the first mechanism, a specific Lys residue of a non-covalently bound acceptor Ub reacts with the thioester bond of the E3-linked Ub (donor Ub), which is accomplished with an E3 monomer and is thus designated the E3 monomer model (Fig. 4B). In the second Ub_2_ formation mechanism, the acceptor Ub covalently binds to the Cys residue on the HECT domain and a specific Lys residue of this acceptor Ub reacts to Ub-G76 on the E2–Ub thiol ester. This is accomplished with an E2/E3 heterodimer and is designated the E2/E3 heterodimer model (Fig. 4B) (16). Wang et al. suggested that the E2/E3 heterodimer model was the most likely mechanism of free Ub_2_ formation by E6AP HECT domain (16). According to the third proposed mechanism, the acceptor Ub is covalently bound to the Cys residue on the HECT domain, and a specific Lys residue of this acceptor Ub attacks the Ub-G76 bond on the second molecule of the HECT-Ub thiolester. This is accomplished with two E3s and therefore this mechanism is designated the E3 homodimer model (Fig. 4B). We first examined the possibility of the E3 monomer model by using Ub_74_ lacking the C-terminus Gly-Gly dipeptide of wild type Ub (Ub_76_). Ub_74_ is not activated by E1 and cannot bind via a thioester bond on HECT, but functions as an acceptor Ub. The E6AP-HECT was preincubated with low concentrations of Ub_76_ to allow Ub_76_ to migrate onto HECT Cys, and then high concentrations of Ub_74_ were added. In E6AP^HECT_WT^, migration of Ub_76_ to HECT Cys was observed, but Ub_2_ formation did not occur (Fig. 4C). This result indicates that E6AP^HECT_WT^ does not form Ub_2_ via the E3 monomer model mechanism, which is consistent with previous reports (16). E6AP^HECT_PPPP^ also did not show Ub_2_ formation, suggesting that E6AP^HECT_PPPP^ froms Ub_2_ model mechanism different from the E3 monomer model (Fig. 4C). We next examined the possibility of Ub_2_ chain formation by E6AP^HECT_PPPP^ according to the E2/E3 heterodimer model. The dependence of E2 concentration on the increasing formation efficiency of the Ub_2_ chain is the basis for the E2/E3 heterodimer model (16). Therefore, we examined the effect of E2 concentration on the free Ub_2_ chain formation efficiency of E6AP^HECT_PPPP^ (Fig. 5A and 5B). Formation of free Ub_2_ increased as the reaction time increased. However, the E2 concentration dependence was not particularly strong over the entire reaction time (Fig. 5A). We also investigated the E3 concentration dependence of the free Ub_2_ chain formation of E6AP^HECT_PPPP^. The free Ub_2_ formation efficiency increased with an increase in E3 concentration (Fig. 5B). These results indicate that the free Ub_2_ formation efficiency more strongly depends on the E3 concentration than on the E2 concentration. The strong dependence of Ub_2_ formation on E3 concentration suggests that Ub_2_ may be synthesized consistent with the E3 homodimer model. However, in this free Ub_2_ formation assay, recycling of E2-Ub occurred in the presence of E1 and Mg/ATP. Under these conditions, Ub_2_ can be formed via both E2/E3 heterodimer and E3 homodimer models throughout the reaction time. To prove Ub_2_ formation mechanism of E6AP^HECT_PPPP^ by E3 homodimer model, we added EDTA after preincubation to prevent E2–Ub recycling by E1 and Mg/ATP. Ten minutes after starting the reaction, a decrease in E2-Ub was observed at 10 min, and no E2-Ub was observed at reaction times longer than 30 min (Fig. 5C). It was indicated that most of the Ub moved from E2 to E6AP HECT domain at 30 min. However, the free Ub_2_ formation still increased after 30 min, even though almost all the E2 existed without carrying Ub (Fig. 5C). The results suggest that the free Ub_2_ chain is formed by E6AP^HECT_PPPP^-Ub molecule, thereby supporting the E3 homodimer model in the case of E6AP^HECT_PPPP^.

**Fig. 4.**
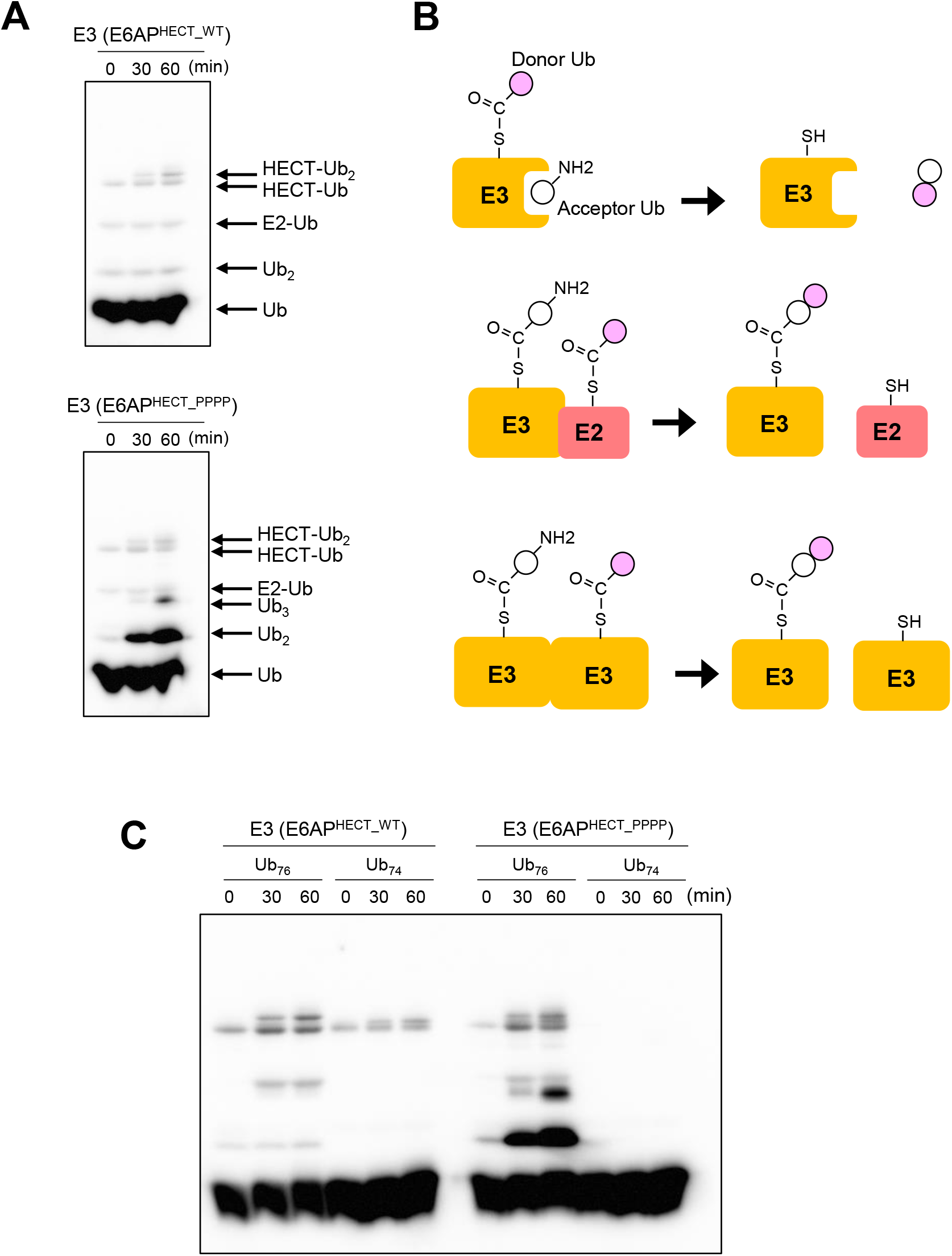
Free Ub_2_ formation by E6AP^HECT_WT^ or UE6AP^HECT_PPPP^. (A) Free Ub_2_ formation by E6AP^HECT_WT^ (upper panel) and E6AP^HECT_PPPP^ (lower panel) was investigated by immunoblot analysis using an anti-Ub antibody. To form E2-Ub thioester complexes, 0.1 μM UBE1 (E1), 6 μM UbcH7 (E2), and 90 μM Ub were pre-incubated for 1 h at 37°C in reaction buffer. After pre-incubation, the reaction mixture was incubated for 10 min at 25°C. Then, 3 μM E6AP^HECT_WT^ or E6AP^HECT_PPPP^ was added to the reaction solutions and reacted for 0–60 min at 25°C. Each lane shows ubiquitinated products corresponding to the indicated reaction time. Arrows indicate the running positions of Ub and the respective Ub thioester complexes. (B) Schematic model of three proposed Ub_2_ formation mechanisms. Top; E3 monomer model, middle; E2/E3 heterodimer model, bottom; E3 homodimer model. The acceptor Ub and donor Ub show in white circle and black circle, respectively. (C) Ub_2_ formation assay by using of Ub_74_ lacking the C-terminus Gly-Gly dipeptide of wild type Ub (Ub_76_). To form E2-Ub thioester complexes, 0.1 μM UBE1 (E1), 6 μM UbcH7 (E2), and low concentration (3 μM) of Ub_76_ were pre-incubated for 1 h at 37°C in reaction buffer. After pre-incubation, the reaction mixture was incubated for 10 min at 25 °C. 3 μM E6AP^HECT_WT^ or E6AP^HECT_PPPP^ was further added and incubate for 10 min at 25 °C to charge Ub_76_ on 3 μM E6AP^HECT_WT^ or E6AP^HECT_PPPP^. Then 90 μM Ub_74_ added to the mixture and reacted for 0–60 min at 25°C.

**Fig. 5.**
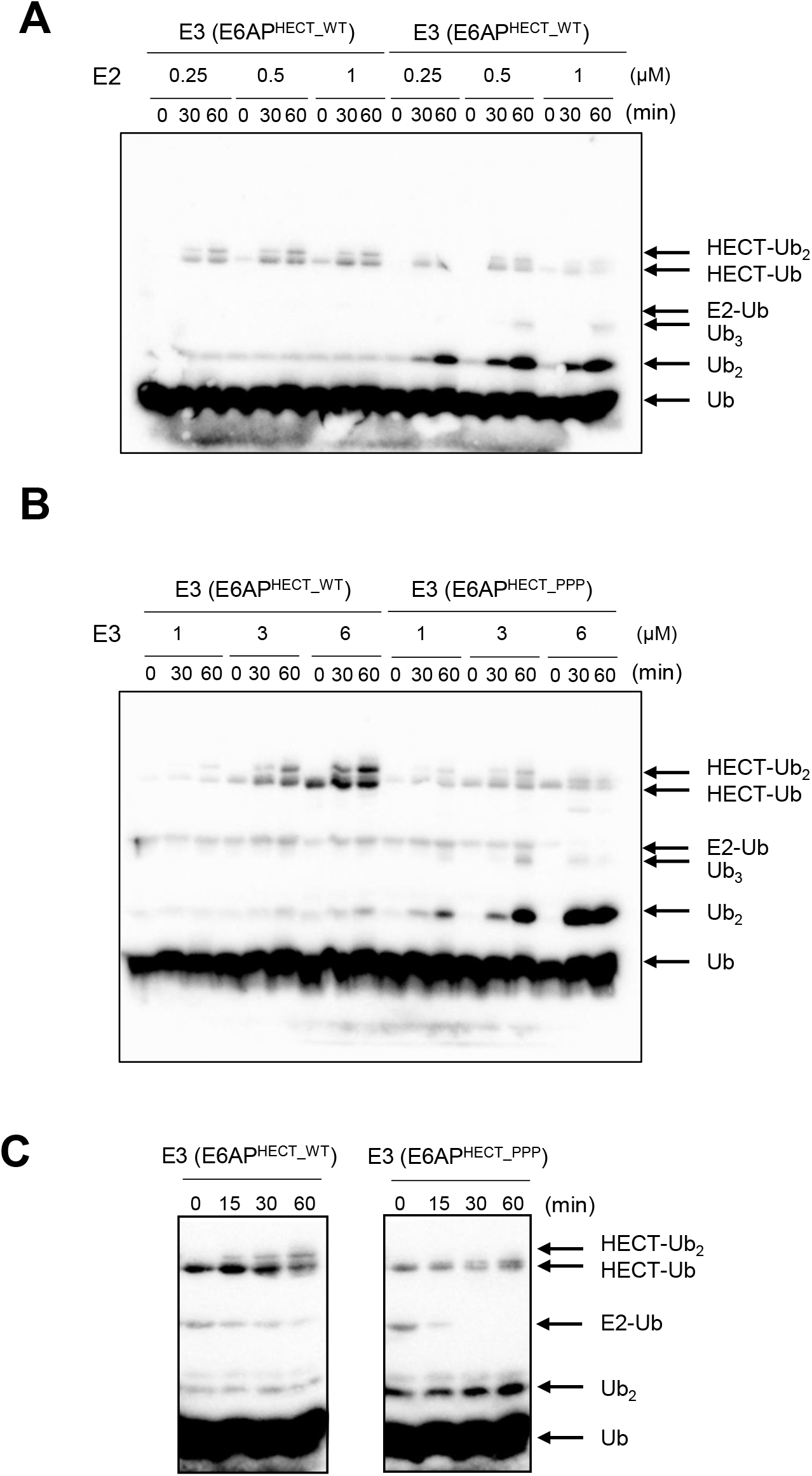
Investigation of free Ub_2_ formation by E6AP^HECT_PPPP^ at different concentration of E2 or E3. (A) Ub_2_ formation assay by at different concentration of E2. 0.1 μM UBE1 (E1), different concentrations of UbcH7 (E2) (1, 3, 6 μM) and 90 μM Ub were pre-incubated for 1 h at 37°C in reaction buffer. After preincubation, the reaction mixture was incubated for 10 min at 25°C. Then, 3 μM of E6AP^HECT_PPPP^ was added to the pre-incubated mixture and reacted for 0–60 min at 25°C. Reaction products were analyzed by immunoblot analysis with anti-Ub antibody. Each lane shows the ubiquitinated products corresponding to the indicated reaction periods. (B) Ub_2_ formation assay by at different concentration of E3. UBE1 (E1), UbcH7 (E2) and Ub were pre-incubated for 1 h at 37°C in reaction buffer. After pre-incubation, the reaction mixture was incubated for 10 min at 25°C. Then, the indicated concentrations of E6AP^HECT_PPPP^ (1, 3, 6 μM) were added to the pre-incubated mixture and reacted for 0–60 min at 25°C. (C) Ub_2_ formation assay without recycling of E2-Ub thioester complex. 0.1 μM UBE1 (E1), 6 μM UbcH7 (E2), and 90 μM Ub were preincubated for 1 h at 37°C in reaction buffer. After pre-incubation, the reaction mixture was incubated with 100 mM EDTA for 10 min at 25°C. Then, 3 μM E6AP^HECT_WT^ or E6AP^HECT_PPPP^ was added to the reaction solutions and reacted for 0–60 min at 25°C.

## Discussion

In this study, we demonstrate the structural dynamics of the E6AP HECT domain in real time. To date, information has only been available from crystal structure analyses (5, 13, 14). Although it was difficult to distinguish the structural state between the reversed T-shape and catalytic conformation, we directly revealed that E6AP^HECT_WT^ adopts each conformational state (L-shape, reversed T-shape or catalytic conformation) with or without Ub. We also revealed the structural state of the hinge loop flexibility-restricted HECT mutant (E6AP^HECT_PPPP^), which has a stable reversed T-shape or catalytic conformation state. Notably, the hinge loop of the HECT domain affects the free Ub_2_ chain synthesis mechanism. The E6AP HECT domain synthesizes a free Ub_2_ chain by the E2/E3 heterodimer model (16). However, in the case of E6AP^HECT_PPPP^, we conclude that the free Ub_2_ chain was mainly formed by the E3 homodimer model with much higher efficiency, though we could not observe dimer formation of E6AT^HECT_PPPP^ in the HS-AFM observation (Fig. 3). We also could not prepare stable E6AT^HECT_PPPP^-Ub by gel filtration. This may be due to the fact that Ub on E6AT^HECT_PPPP^ quickly converts to Ub_2_ and dissociates from E6AT^HECT_PPPP^. However, we sometimes could observe an oligomer structure of E6AT^HECT_WT^-Ub that was not observed in the case of E6AT^HECT_WT^ (Fig. S3). In addition, it is known that full-length E6AP forms an oligomer, and the oligomer shows the maximum activity of Ub delivery to the target protein (20). The possibility that dimer or oligomer formation of E6AP HECT domain transiently occurs only when the E6AP HECT domain contains Ub remains. In addition, our results also suggest that the binding affinity of each E6AP HECT domain containing Ub becomes higher when E6AP HECT domains take reversed T-shape or catalytic conformation state. Considering our results of HS-AFM observation and biochemical experiments, we conclude that the E3 homodimer model presents the most likely explanation of Ub_2_ formation by E6AP^HECT_PPPP^.

There are several interpretations for the importance of the hinge loop that can be made from these results (Fig. 6). First, free Ub_2_ seems to be formed mainly by E2/E3 heterodimer model. When HECT adopts an L-shaped structure after receiving Ub, the E2 binding site on HECT becomes empty, and other E2-Ub binds there. However, since the C-lobe of the L-shape is far from E2-Ub, it is disadvantageous for Ub_2_ formation. In fact, we observed that E6AP^HECT_WT^ had a lower efficiency of free Ub_2_ formation than E6AP^HECT_PPPP^. However, Ub_2_ formation mechanism depends on how the hinge loop works. In the reversed T-shape or catalytic conformation state after receiving Ub, the binding of E2-Ub to the HECT domain is prevented; instead, dimer formation of E3 is increased, and free Ub_2_ is formed efficiently because of the close position of Ubs on each E3 (Fig. 6). Thus, the hinge loop is not only important for the transfer of Ub to the target protein, but also for free Ub_2_ chain formation mechanism. The formation efficiency of the free Ub_2_ and longer poly-Ub chain of KIAA 10, a type-E3 HECT, is much higher than that of the E6AP HECT domain (16). Direct observation of the structural dynamics of KIAA 10 should provide a better understanding of the relationship between Ub_2_ chain formation and the hinge loop.

**Fig. 6.**
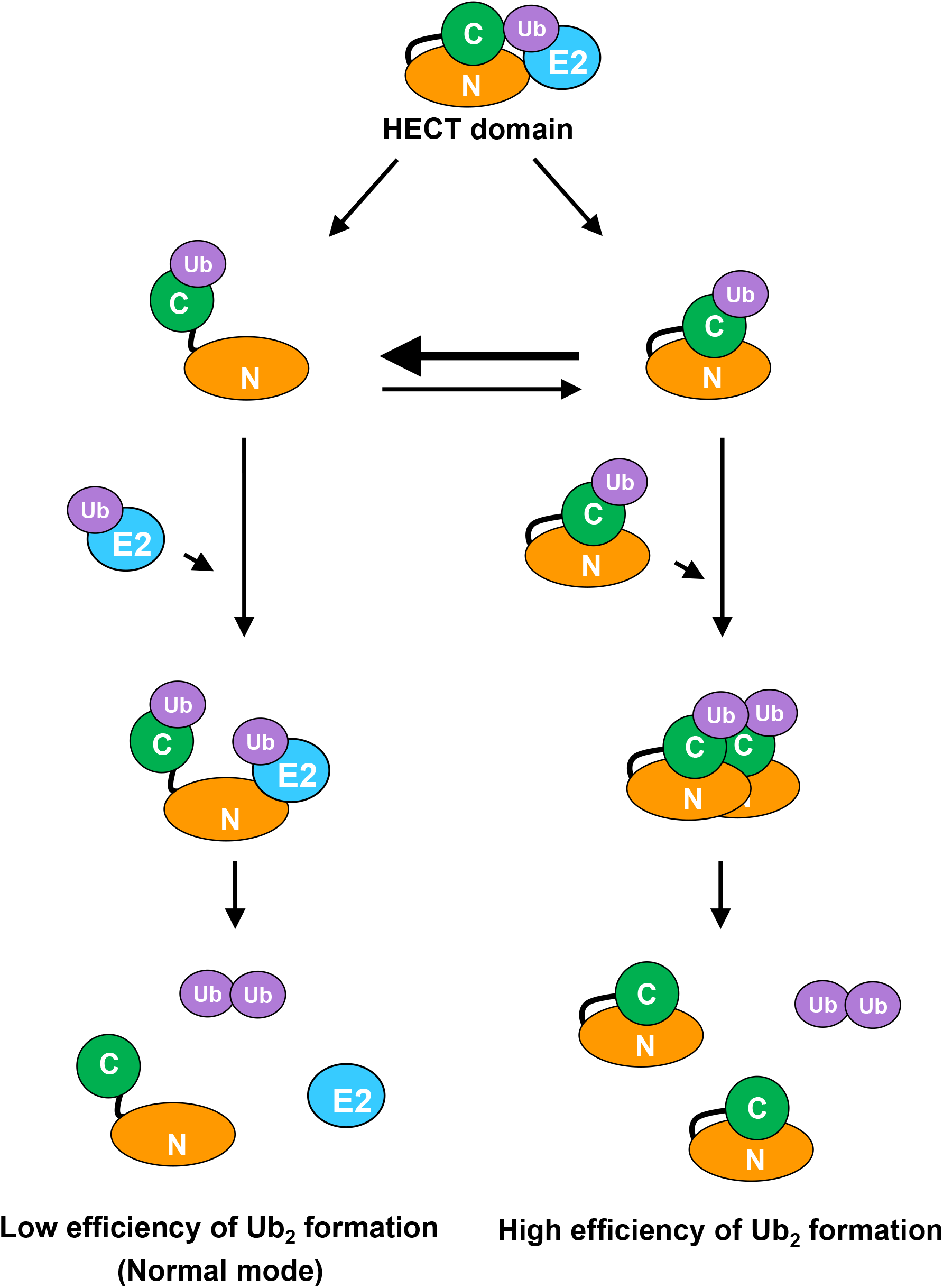
Schematic illustration of free Ub_2_ formation efficiency by E6AP HECT domain depending on its structural and association state. Two Ub_2_ formation models depending on the structural and association states of the E6AP HECT domain. The left pathway depicts the low efficiency of Ub_2_ formation. When the HECT domain adopts an L-shaped structure after receiving Ub, the E2 binding site on the HECT domain becomes empty, and the other E2-Ub binds to the binding site. Ub_2_ is then formed with low efficiency because the C-lobe of the L-shape is far from E2-Ub. The right pathway depicts the high efficiency of Ub_2_ formation. When the HECT domain adopts the T-shape or conformational state structure after receiving Ub, the formation of HECT domain dimers occurs and Ub_2_ is formed efficiently because of the close position of Ubs on each HECT domain.

Taken together, our results indicate that the hinge loop flexibility of the HECT domain has a great influence not only in transferring Ub to target proteins, but also in the formation mechanism of the free Ub_2_ chain. These results also predict that the hinge loop functions in efficiently transferring Ub from E3 to target proteins as well as in avoiding wasteful Ub_2_ chain formation *in vivo*. Although the physiological relevance of the free Ub_2_ chain remains unclear, in the linear Ub assembly complex RING-type E3, the binding of Ub_2_, which is not related to Ub transfer to the target protein, greatly increases the Ub transfer activity (21). Thus, the effects of free Ub_2_ on the activity of various E3s and related intracellular processes need to be further investigated.

## Materials & Methods

### Cloning, expression, and purification of the wild-type E6AP HECT domain (E6AP^HECT_WT^) and hinge loop flexibility-restricted mutant (E6AP^HECT_PPPP^)

DNA fragments coding the C-terminal 350 amino acids of E6AP were synthesized as a codon-optimized artificial gene (GenScript, Piscataway, NJ, USA). The products were subsequently ligated into the pET28a vector with NdeI and BamHI restriction enzymes and designated E6AP^HECT_WT^. *Escherichia coli* BL21 (DE3) harboring E6AP^HECT_WT^ expression plasmids were cultured in 2×YT medium containing 40 μg/mL kanamycin at 25°C to an optical density at 600 nm (OD_600_) of 0.6–0.8. Protein expression was then induced by the addition of 1 mM isopropyl β-d-1-thiogalactopyranoside for 4 h at 25°C. Cells were collected by centrifugation, suspended in 20 mM MOPS-KOH (pH 7.0), 150 mM NaCl, and 5 mM 2-Mercaptoethanol (2-ME), and disrupted using a French press. The supernatant obtained after lysate centrifugation (17,800 × g, 15 min, 4°C) was applied to a Toyopearl DEAE-650M column (Tosoh, Tokyo, Japan) equilibrated with the same buffer. The column was washed with 200 mL of the same buffer solution. The washed fraction was applied to a Ni-NTA Superflow column (QIAGEN, Hilden, Germany) equilibrated with 20 mM MOPS-KOH (pH 7.0), 300 mM NaCl, and 5 mM 2-ME (pH 7.0). The column was washed with 20 mM MOPS-KOH (pH 7.0), 300 mM NaCl, 100 mM imidazole, and 5 mM 2-ME. His-E6AP^HECT_WT^ was eluted with 20 mM MOPS-KOH (pH 7.0), 300 mM NaCl, 200 mM imidazole, and 5 mM 2-ME. The elution fraction was concentrated using Amicon® Ultra Centrifugal Filters with a molecular weight cut-off (MWCO of 30,000 (Millipore, Billerica, MA, USA). The concentrated sample was applied to a Superdex 75 gel filtration chromatography column equilibrated with 50 mM HEPES-NaOH (pH 7.0), 100 mM NaCl, and 1 mM dithiothreitol (DTT) at a flow rate of 0.5 mL/min. Elution was monitored at 280 nm. The concentration of the eluted E6AP^HECT_WT^ was quantitated using the Bradford method (Bio-Rad, Hercules, CA, US), and the product was stored at −80°C until use. To introduce a mutation that restricts the flexibility of the hinge loop, the GSRN sequence in E6AP^HECT_Wt^ was substituted with PPPP using the KOD-Plus-Mutagenesis Kit (TOYOBO, Osaka, Japan) using the mutation primers 5-CTGTCCGCCACCACCGCTGGATTTCCAGGC −3 and 5’-ATCAGCAGTTCGATTTCTTCGGGACGAAAG −3’. The generated mutant was designated E6AP^HECT_PPPP^. The E6AP^HECT_PPPP^ protein was expressed and purified as described in the E6AP^HECT_WT^ preparation.

### Preparations of Ub, E1 (UBE1), and E2 (UBCH7)

Cloning, expression, and purification of Ub, E1 (UBE1), and E2 (UBCH7) was performed as previously described (22).

### Free Ub_2_ formation assays

To form E2-Ub thioester complexes, 0.1 μM E1 (UBE1) and 6 μM E2 (UBCH7) were pre-incubated with 90 μM Ub for 1 h at 37°C in reaction buffer (25 mM HEPES-NaOH (pH7.0) 100 mM NaCl, 10 mM ATP, 10 mM MgCl2). After pre-incubation, the reaction mixture was incubated for 10 min at 25°C. Then, E6AP^HECT_WT^ or E6AP^HECT_PPPP^ (3 μM each) was added to the reaction solutions and reacted for 0–60 min at 25°C. Reactions were quenched by adding 5× SDS-PAGE loading buffer lacking DTT [0.25 M Tris-HCl (pH 6.8), 50% (w/v) glycerol, 10% (w/v) SDS, 0.25% (w/v) bromophenol blue]. Whole reaction mixtures were separated on 12% wide-range gels (Nacalai Tesque, Kyoto, Japan) and detected by western blot analysis using anti-Ub antibody. To perform Ub_2_ assay in the absence of E2-Ub recycling, the preincubated sample described above was incubated with 100 mM EDTA 10 min at 25°C before adding of E6AP^HECT_WT^ or E6AP^HECT_PPPP^.

### Preparation of E6AP^HEC_WT^-Ub thioester complexes

For isolation of E6AP^HECT_WT^-Ub thioester complexes, 100 nM E1 and 4.5 μM E2 were pre-incubated with 9 μM Ub for 1 h at 37°C in reaction buffer (25 mM HEPES-NaOH (pH7.0) 100 mM NaCl, 10 mM ATP and 10 mM MgCl2). After pre-incubation, the reaction mixture was incubated for 10 min at 25°C. E6AP^HECT_WT^ (3 μM) was added to the reaction solutions and reacted for 15 min at 25°C to form the E6AP^HECT^-Ub complexes. The ubiquitinated reaction mixture was applied to a Ni-NTA super flow column equilibrated with 20 mM MOPS-KOH and 300 mM NaCl (pH 7.0). After washing with 20 mM MOPS-KOH, 300 mM NaCl, and 50 mM imidazole-HCl (pH 7.0), the E6AP^HECT_WT^-Ub complex was eluted and isolated using 20 mM MOPS-KOH, 300 mM NaCl, and 200 mM imidazole-HCl (pH 7.0). The concentration of the eluted E6AP^HECT_WT^-Ub was quantitated using the Bradford method, and the product was stored at −80°C until use.

### HS-AFM observation of the E6AP^HECT_WT^, E6AP^HECT_WT^-Ub complex and E6AP^HECT_PPPP^

AFM imaging of the E6AP HECT domains in solution was performed using a laboratory-built HS-AFM setup (23, 24). Two microliters of a 15 nM E6AP HECT domain sample were placed on a mica substrate and incubated for 10 min at room temperature (22-28°C). Unattached molecules were removed by washing with observation buffer (50 mM HEPES-NaOH (pH 7.0) 100 mM NaCl). Imaging was carried out in tapping mode using small cantilevers (BL-AC10DS-A2 or custom-made BL-AC7DS-KU4; Olympus, Tokyo, Japan). The cantilever-free oscillation amplitude was approximately 1.5 nm, and the set-point amplitude was 80-90% of the free oscillation amplitude. The imaging rate, scan size, and feedback parameters were optimized to enable visualization using a minimum tip force. AFM data analysis were performed using “Falcon viewer” custom software.

### Automatized fitting of PDB structures into experimental AFM image

Simulation AFM and automatized fitting was performed within the BioAFMviewer interactive software interface (25). Simulated scanning was based on the non-elastic collisions of a cone-shaped scanning tip with a rigid-sphere atomic model of the PDB protein structure (see (25) for details). The automatized fitting procedure identified the optimal match of simulated AFM image and the target experimental HS-AFM image (26). Optimal fitting results were obtained with a tip having 3nm probe-sphere radius and a cone half-angle of 10°.

## Acknowledgments

This work was supported by a KAKENHI grant for Research on Innovative Area (#25112507 to H.K.), the World Premier International Research Center Initiative (WPI), MEXT, Japan, and the Kanazawa University CHOZEN project.

## Author contributions

H.K., K.T., and I.M. designed the research; F.K. and I.M. performed the biological experiments; H.K., K.T., and I.M. performed the HS-AFM observations; H.K., K.T., I.M. analyzed the data; H.F. performed simulation AFM; and H.K., K.T., I.M., N.K., and T.A. wrote the paper with input from all authors.

The authors declare no conflict of interest.

---

**Fig. S1.**
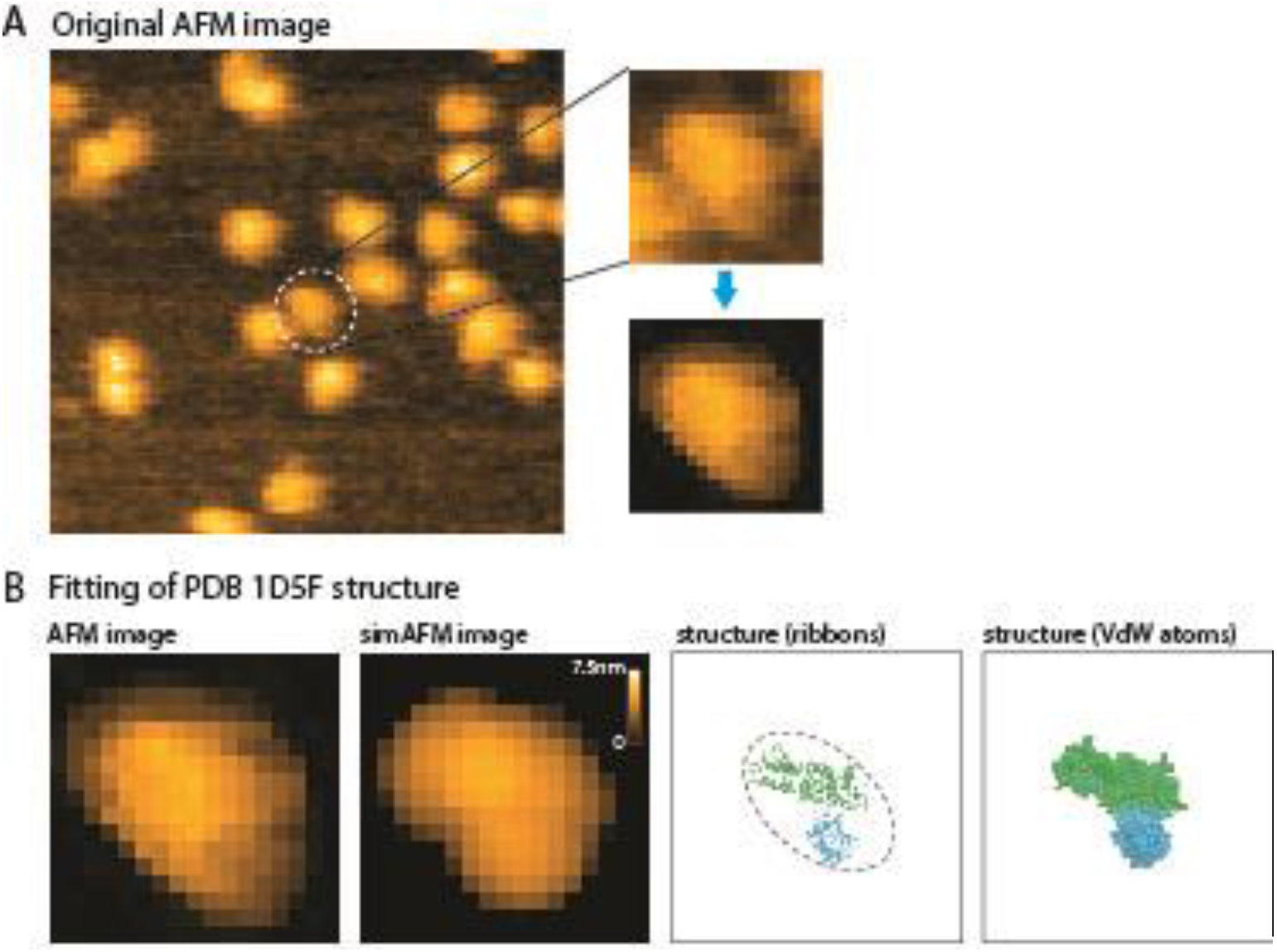
Automatized fitting of PDB structure 1D5F into experimental AFM image. A) Original HS-AFM image, a selected section (upper right) and the cleaned version used for fitting (bottom right). B) HS-AFM image and the best-match simulation AFM image obtained from structural fitting (left). Their similarity is quantified by an image correlation coefficient of 0.96. The identified underlying molecular structure is shown on the right side in ribbon and atomic representations.

**Fig. S2.**
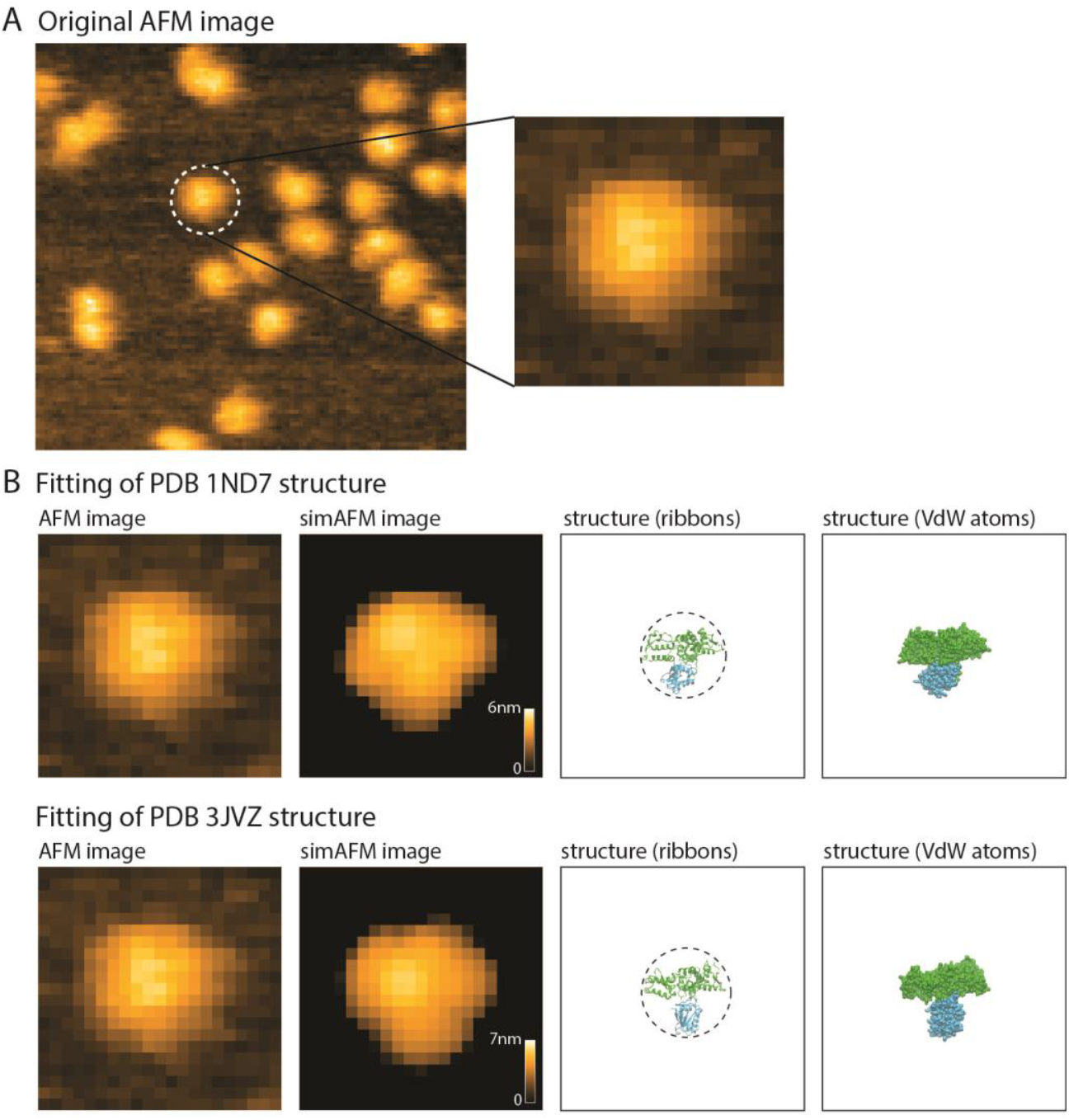
Automatized fitting of PDB structures 1ND7 and 3JVZ into experimental AFM image. A) Original HS-AFM image and a selected section used for fitting (right). B) Upper panel: HS-AFM image and the best match simulation AFM image obtained from structural fitting of PDB 1ND7 (left). Their similarity is quantified by an image correlation coefficient of 0.95. The identified underlying molecular structure is shown on the right side in ribbon and atomic representations. Bottom panel: HS-AFM image and the best match simulation AFM image obtained from structural fitting of PDB 3JVZ (left). Their similarity is quantified by an image correlation coefficient of 0.94. The identified underlying molecular structure is shown on the right side in ribbon and atomic representations.

**Fig. S3.**
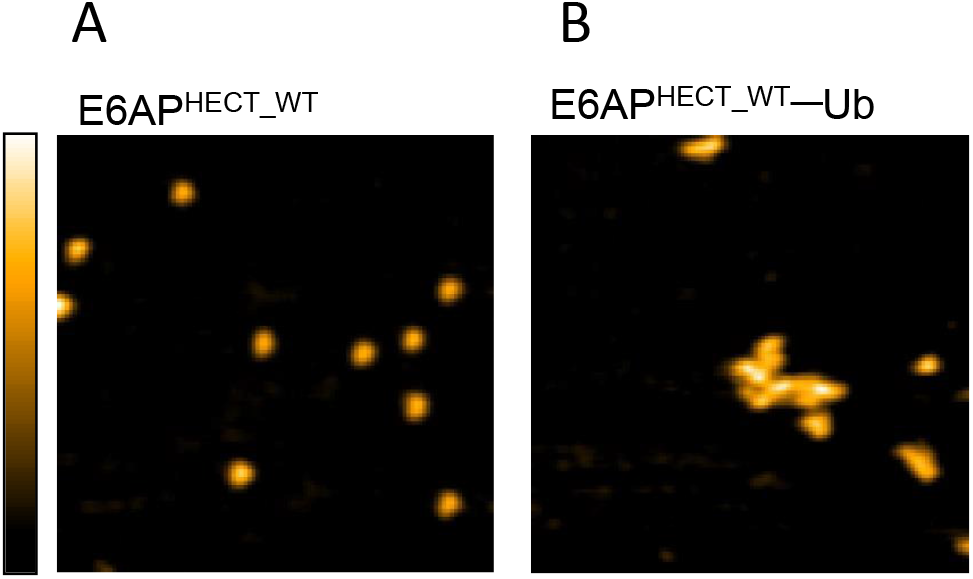
AFM images of E6AP^HECT_WT^ in the absence and presence of Ub. AFM image of E6AP^HECT_WT^ (A) and E6AP^HECT_WT^–Ub (B). E6AP^HECT_WT^–Ub was prepared as described in material and methods. Imaging rate, 1.67 frames/s (fps); Scan range, 200 nm × 200 nm (120pixel × 120pixel); Z-scale, 4 nm.

